# Quantitatively Partitioning Microbial Genomic Traits among Taxonomic Ranks across the Microbial Tree of Life

**DOI:** 10.1101/520973

**Authors:** Taylor M. Royalty, Andrew D. Steen

## Abstract

Widely used microbial taxonomies, such as the NCBI taxonomy, are based on a combination of sequence homology among conserved genes and historically accepted taxonomies, which were developed based on observable traits such as morphology and physiology. A recently-proposed alternative taxonomy, the Genome Taxonomy Database (GTDB), incorporates only sequence homology of conserved genes and attempts to partition taxonomic ranks such that each rank implies the same amount of evolutionary distance, regardless of its position on the phylogenetic tree. This provides the first opportunity to completely separate taxonomy from traits, and therefore to quantify how taxonomic rank corresponds to traits across the microbial tree of life. We quantified the enrichment of clusters of orthologous gene functional categories (COG-FCs) as a proxy for traits within the lineages of 13,735 cultured and uncultured microbial lineages from a custom-curated genome database. On average, 41.4% of the variation in COG-FC enrichment is explained by taxonomic rank, with domain, phylum, class, order, family, and genus explaining, on average, 3.2%, 14.6%, 4.1%, 9.2%, 4.8%, and 5.5% of the variance, respectively (*p*<0.001 for all). To our knowledge, this is the first work to quantify the variance in metabolic potential contributed by individual taxonomic ranks. A qualitative comparison between the COG-FC enrichments and genus-level phylogenies, generated from published concatenated protein sequence alignments, further supports the idea that metabolic potential is taxonomically coherent at higher taxonomic ranks. The quantitative analyses presented here constrain the integral relationship between diversification of microbial lineages and the metabolisms which they host.

**Importance:** Recently there has been great progress in defining a complete taxonomy of bacteria and archaea, which has been enabled by improvements in DNA sequencing technology and new bioinformatic techniques. A new, algorithmically-defined microbial tree of life describes those linkages relying solely on genetic data, which raises the question of how microbial traits relate to taxonomy. Here, we adopted cluster of orthologous group functional categories as a scheme to describe the genomic contents of microbes, which can be applied to any microbial lineage for which genomes are available. This simple approach allows quantitative comparisons between microbial genomes with different gene composition from across the microbial tree of life. Our observations demonstrate statistically significant patterns in cluster of orthologous group functional categories at the taxonomic levels spanning from domain to genus.

## Introduction

The relationship between microbial taxonomy and function is a longstanding problem in microbiology (1–3). Prior to the identification of the 16S rRNA gene as a taxonomic marker, microbial phylogenetic relationships were defined by traits such as morphology, behavior, and metabolic capacity. Cheap DNA sequencing has provided the ability to fortify those phenotype-based taxonomies with quantitative determinations of differences between marker genes, but canonical taxonomies such as the NCBI taxonomy continue to “reflect the current consensus in the systematic literature,” which ultimately derives from trait-based taxonomies (4). Recently, Parks et al. (5) formalized the genome taxonomy database (GTDB), a phylogeny in which taxonomic ranks are defined by “relative evolutionary divergence” in order to create taxonomic ranks that have uniform evolutionary meaning across the microbial tree of life (5). This approach removes phenotype or traits entirely from taxonomic assignment. Thus, an investigation of the relationship between traits and phylogeny has not been possible until the recent publication of a microbial tree of life that is based solely on evolutionary distance.

Comparing phenotypic characteristics of microorganisms across the tree of life is not currently possible, because most organisms and lineages currently lack cultured representatives (6, 7). We therefore used the abundance of different Clusters of Orthologous Groups (COGs) in microbial genomes, a proxy for phenotype which is available for all microorganisms for which genomes are available. Clusters of orthologous groups (COGs) are a classification scheme that defines protein domains based on groups of proteins sharing high sequence homology (8). More than ~5,700 COGs have been identified to date. COGs are placed into one of 25 metabolic functional categories (COG-FCs), which represents a generalized metabolic function. Our analyses quantify the degree to which taxonomic rank (genus through domain) predicts the COG-FC content of genomes, and illustrate which lineages are relatively enriched or depleted in specific COG-FCs. These analyses constitute a step towards being able to probabilistically predict the metabolic or functional similarity of microbes given their taxonomic classification.

## Results

The genomes analyzed in this work were compiled from a variety of different sources, including RefSeq v92, JGI IMG/M, and Genbank, in order to include genomes created using diverse sequencing and assembly techniques. The integration of RefSeq v92, JGI IMG/M, and Genbank databases resulted in a total of 119,852 genomes within the custom-curated database. Raw data, GTDB taxonomy, and associated accessions are provided in Dataset S1, which is explained in more detail in Supplement 3, available here: https://www.dropbox.com/sh/rnm6ount2aqkmvn/AACGgZhdrA0fSYnD0mNtIpaxa?dl=0]. Of these genomes, we included only those that satisfied a set of criteria designed to ensure that each genus contained enough genomes to allow statistically robust analysis (see Methods). This resulted in a set of 13,735 lineages, representing 22 bacterial phyla and 4 archaeal phyla, of which 67% have been grown in culture (Table 1).

**Table 1.** A summary of the custom-curated genome database used in this work.

| Unique Domains | Unique Phyla | Unique Classes | Unique Orders | Unique Families | Unique Genera | Unique Lineages | Cultured Lineages | Uncultured Lineages |
| --- | --- | --- | --- | --- | --- | --- | --- | --- |
| Bacteria | Actinobacteriota | 3 | 9 | 22 | 50 | 2286 | 2115 | 171 |
|  | Bacteroidota | 3 | 7 | 19 | 50 | 1606 | 741 | 865 |
|  | Campylobacterota | 1 | 1 | 6 | 8 | 270 | 203 | 67 |
|  | Cyanobacteriota | 2 | 3 | 4 | 7 | 119 | 84 | 35 |
|  | Deinococcota | 1 | 1 | 2 | 2 | 44 | 44 | 0 |
|  | Desulfobacterota | 2 | 2 | 2 | 4 | 43 | 23 | 20 |
|  | Elusimicrobiota | 1 | 1 | 1 | 1 | 22 | 0 | 22 |
|  | Fibrobacterota | 1 | 1 | 1 | 1 | 34 | 22 | 12 |
|  | Firmicutes | 3 | 10 | 23 | 48 | 1543 | 1356 | 187 |
|  | Firmicutes A | 2 | 7 | 10 | 31 | 600 | 304 | 296 |
|  | Firmicutes B | 1 | 1 | 1 | 1 | 22 | 11 | 11 |
|  | Firmicutes C | 1 | 2 | 3 | 4 | 53 | 32 | 21 |
|  | Fusobacteriota | 1 | 1 | 2 | 2 | 40 | 40 | 0 |
|  | Marinisomatota | 1 | 1 | 1 | 1 | 10 | 0 | 10 |
|  | Nitrospirota | 2 | 2 | 2 | 2 | 30 | 6 | 24 |
|  | Nitrospirota A | 1 | 1 | 1 | 1 | 14 | 2 | 12 |
|  | Patescibacteria | 6 | 16 | 26 | 36 | 707 | 0 | 707 |
|  | Proteobacteria | 3 | 25 | 59 | 163 | 5589 | 3952 | 1637 |
|  | Spirochaetota | 3 | 4 | 4 | 6 | 153 | 89 | 64 |
|  | Synergistota | 1 | 1 | 1 | 1 | 19 | 2 | 17 |
|  | Thermotogota | 1 | 1 | 2 | 2 | 23 | 15 | 8 |
|  | Verrucomicrobiota | 2 | 4 | 5 | 7 | 84 | 16 | 68 |
| Archaea | Crenarchaeota | 1 | 1 | 1 | 2 | 68 | 7 | 61 |
|  | Euryarchaeota | 2 | 2 | 2 | 2 | 45 | 26 | 19 |
|  | Halobacterota | 4 | 5 | 7 | 9 | 164 | 97 | 67 |
|  | Thermoplasmatota | 1 | 1 | 2 | 9 | 147 | 0 | 147 |
| Total | 26 | 50 | 110 | 209 | 450 | 13,735 | 9187 | 4548 |

Most predicted open reading frames for most lineages could be assigned to a COG-FC. Across all phyla, an average of 84.3% + 7.8% of open reading frames were assigned to a COG-FC (Fig S1). Genomes of the same phylum tended to group together in an initial principal component analysis (PCA) of raw COG-FC abundance (Fig. 1A). Since this analysis was based on absolute abundance of COG-FCs in genomes, rather than relative abundance, we hypothesized that the relationship between COG-FC abundance and phylum was largely a consequence of genome size, which is phylogenetically conserved (9). Consistent with this possibility, position on PC 1 correlated closely with genome size (R^2^=0.88; Fig 1B). We therefore normalized each COG-FC abundance, for each genome, to a prediction of COG-FC abundance as a function of genome size derived from a generalized additive model (GAM; Fig S2; summary statistics in table S1). Each GAM model was statistically significant (*p* < 0.001), and all but five COG-FCs had deviance explained (analogous to adjusted R^2^) of more than 50%. We interpret analyses of these genome size-normalized data sets as reflecting the enrichment or depletion of COG-FC abundance, relative to that expected for a given genome size. PCA of these COG-FC enrichments showed that species-level lineages still tended to group by phylum, even though the inter-phyla gradients in genome size were no longer apparent (Fig 1C, D).

**Fig 1.**
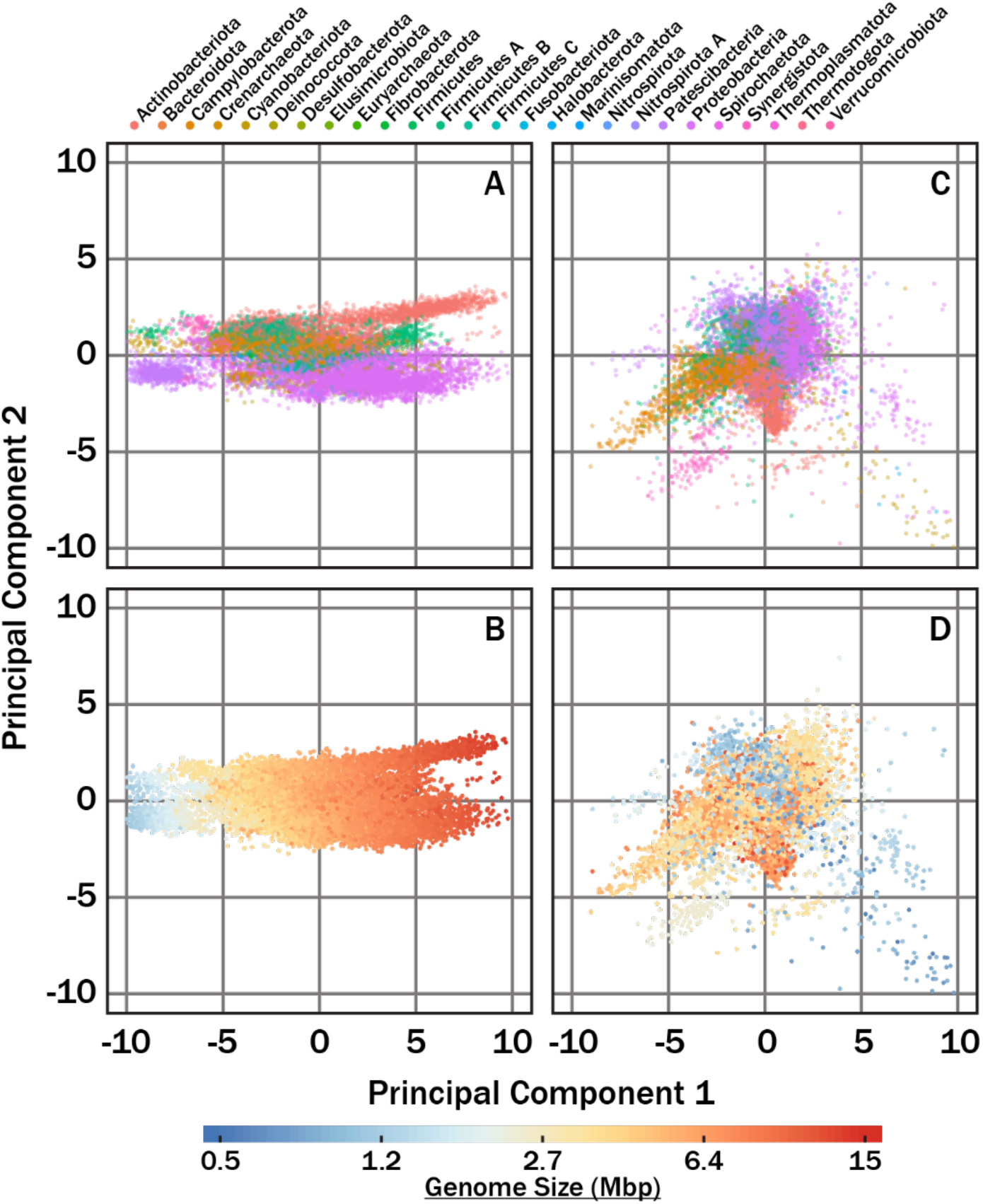
PCA plots of COG-FC abundance (A,B) and enrichment (C,D) of the lineages summarized in Table 1. Individual data points are colored by genome size (A,C) and phylum (B, D). Panels A and B were not normalized by genome size while panels C and D were normalized by genome size. For panels A and B, PC1 explained 71% and PC2 explained 7.0% of variance. For panels C and D, PC1 explained 21% and PC2 explained 16% of variance.

To quantify the degree to which taxonomic rank explains the distribution of COG-FC enrichments among individual genomes, we performed a permutation multivariate ANOVA (PERMANOVA) using the following taxonomic ranks: domain, phylum, class, order, family, and genus, as well as culture-status (cultured versus uncultured lineage). The rank of species was excluded from the analysis as every lineage was unique, and thus, species would explain 100% of the data. Every rank significantly influenced the distribution of COG-FC enrichments (*p* < 0.001), but the fraction of variance that each rank explained differed substantially: phylum explained the most variance (14.6%), followed by order (9.2%), genus (5.5%), family (4.8%), and class (4.1%). Domain explained only 3.1% of variance in COG-FC enrichment, the least of any taxonomic rank. Culture-status was a significant correlate of COG-FC abundance (*p* < 0.001) but had virtually no explanatory power, with variance explained <0.001%. This observation is consistent with no particular COG-FC being systematically enriched or depleted in uncultured microbes relative to cultured microbes.

The variability in COG-FC enrichment across different phyla was explored in addition to mean COG-FC composition for individual phyla (Fig 3). Here, the distance in PCA space between each lineage and the centroid of all lineages in its phyla is a measure of how different the genome is, in terms of COG-FC distribution, from the typical lineage in that phylum. Among all phyla, COG-FC distribution of Crenarchaeota was the most variable, followed by Patescibacteria and Cyanobacterota. The least variable phyla were the Synergistota, Marinisomatota, and Fibrobacterorta, respectively (Fig 3A). We explored the possibility that variance in COG-FC distribution was a function of the number of lineages in the phylum. In other words, did COG-FC content of some genomes simply seem less variable because they had been under-sampled? A plot of the average distance of lineages from their phylum’s centroid versus the number of lineages in the phylum reveals an apparent increase in variability between 10 and roughly 100 genomes, at which point average distance being approaching an asymptote (Fig 3B). Comparison of Akaike Information Criterion (AIC) of a saturating model of the data (Eq. 1), fit using nonlinear least squares, to a linear regression indicated that the saturating model described the relationship substantially better (ΔAIC = 10.5). Coefficient *A* of the saturating model, which represents the value of the asymptote, was estimated to be 0.75 +/− 0.15 (p < 0.001). Coefficient B, which represents how quickly the function approaches the asymptote, was 0.43 +/− 0.30 (p = 0.17). Coefficient C, an offset to handle the fact that all the log-transformed distances have negative values, was −1.63 +/− 0.14 (p < 0.001).

**Fig 2.**
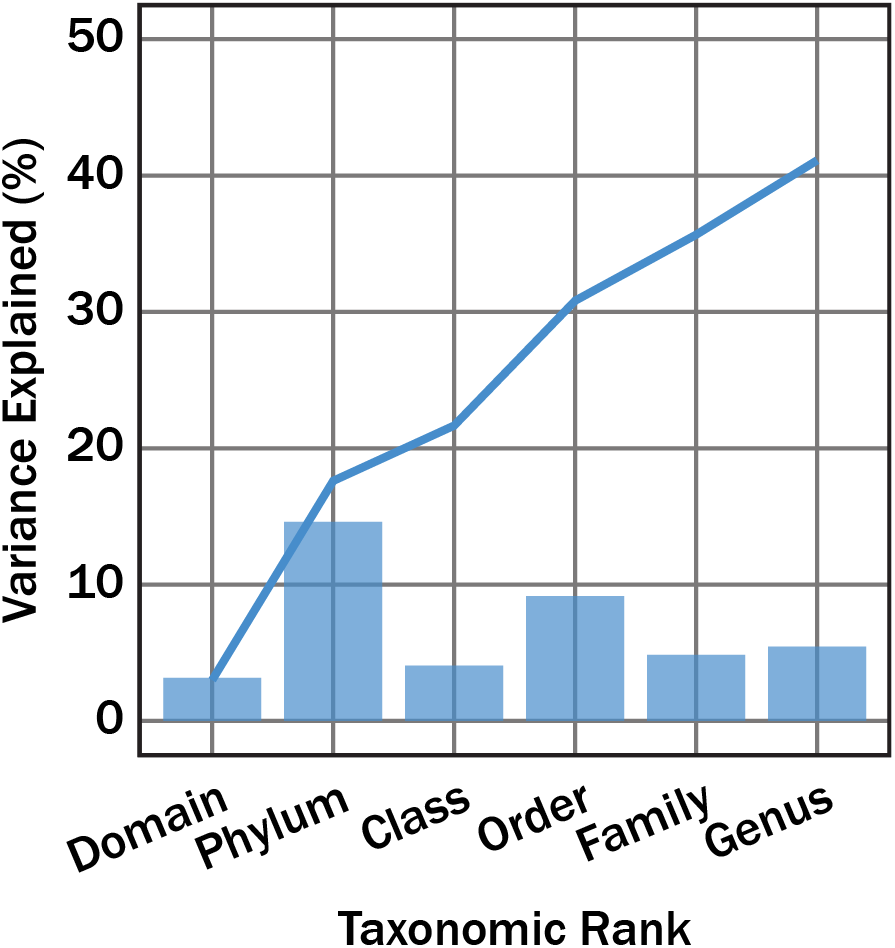
The average variance in COG-FC enrichment explained by different taxonomic ranks (bars) and the cumulative variance explained by taxonomic ranks (lines). All variance explained by taxonomic ranks was significant (*p*<0.001). The *F*-value for domain, phylum, class, order, family, and genus, was 726.0, 128.8, 38.76, 34.4, 11.2, and 5.1, respectively.

**Fig 3.**
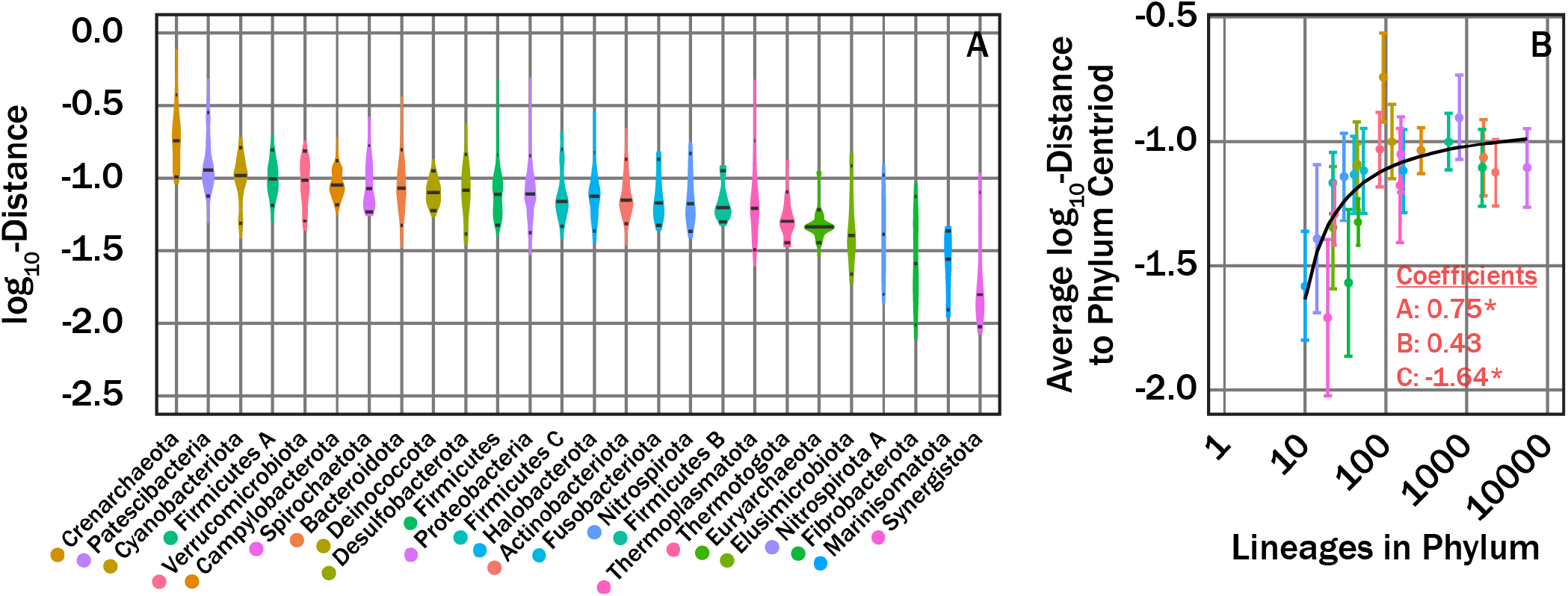
Violin plots showing the distribution in log_10_-distance for lineages from their respective phyla mean COG-FC centroid (A) and average log_10_-distance to phyla mean COG-FC centroid as a function of number of lineages in a given phylum (B). Coefficients in panel B correspond to fit parameters from eq 1. The * symbols denote significance as defined in the text. We note three outliers: the Crenarhaeota are characterized by unusually high diversity of COG-FCs distributions, and the Synergistota and Fibrobacterota are characterized by an unusually low diversity of COG-FC distributions.

Finally, we quantified the average enrichment or depletion of each COG-FC in each genus These values were then visualized on a genus-level phylogenetic tree (Fig 4) built from concatenated ribosomal protein sequences published by Parks et al. (5). Data underlying Fig. 4 are presented in Supplemental Data Set 2. Several notable features appear in COG-FC enrichments at the phylum level. For example, among the four Archaeal phyla represented here, Thermoplasmatota appears unique, with substantial enrichments in cell motility and depletion in every other category. In general, the bacteria appeared more variable at the phylum level than the archaea. The clade consisting of Bacteroidota, Spirochaetota, and Verrucomicrobiota were notably depleted in the less-variable COG-FCs, including energy production and conversion, amino acid transport and metabolism, and carbohydrate transport and metabolism, among others. Another prominent feature is the near-ubiquitous enrichment of cell motility, secondary metabolites biosynthesis, transport, and catabolism, lipid transport and metabolism, and intracellular trafficking, secretion, and vesicular transport COGs in Proteobacteria. A notable dichotomy in the enrichment of RNA processing and modification within the Proteobacteria mirrors the division of the two largest clades within the proteobacteria. Overall, enrichment data qualitatively appears consistent with phylogenetic relationships, albeit, occurring on different taxonomic levels.

**Fig 4.**
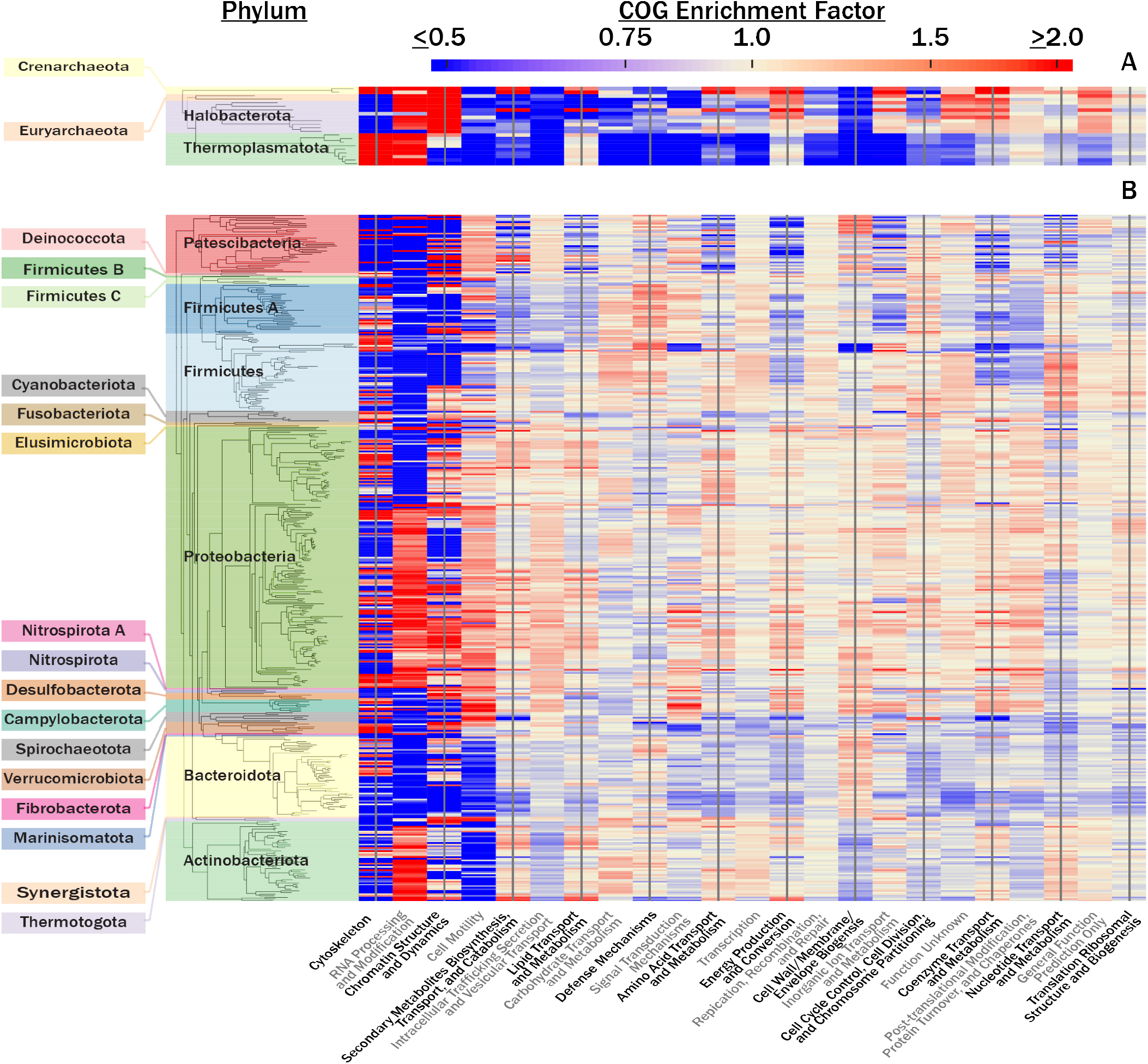
A heat map showing the average COG-FC enrichment for all archaeal (A) and bacterial (B) genera. Categories were arranged from left to right along the x axis in order of decreasing total variance in enrichment across all lineages. Clades were organized along the y axis using phylogenetic relatedness based on the reported concatenated protein sequence alignments in Parks et al. (1).

To gain a sense of “notable” COG-FCs associated with different phyla, we calculated the mean COG-FC across all lineages in a given phyla and compared these values against the 85^th^ and 15^th^ percentiles for all lineages in our custom-curated database. All COG-FCs which were significantly (*p* < 0.05; based on a 10^5^-iteration bootstrap analysis) greater or less than the 85^th^ and 15^th^ percentiles, respectively, are shown in Table 2. Each archaeal phylum was enriched or depleted three-to-nine COG147 FCs, whereas most bdacteria phyla were enriched or depleted in in 3-4 COG-FCs. A few exceptions arose, such as Fibrobacterota was deplete in 8 COG-FCs, Nitrospirota A was enriched in 4 and depleted in 5, and Proteobacteria was the only phylum not heavily enriched or depleted in any COG-FCs. Enrichment data, along with associated GTDB taxonomic assignments, used for generating Fig 4 is available in Dataset S2.

**Table 2.**
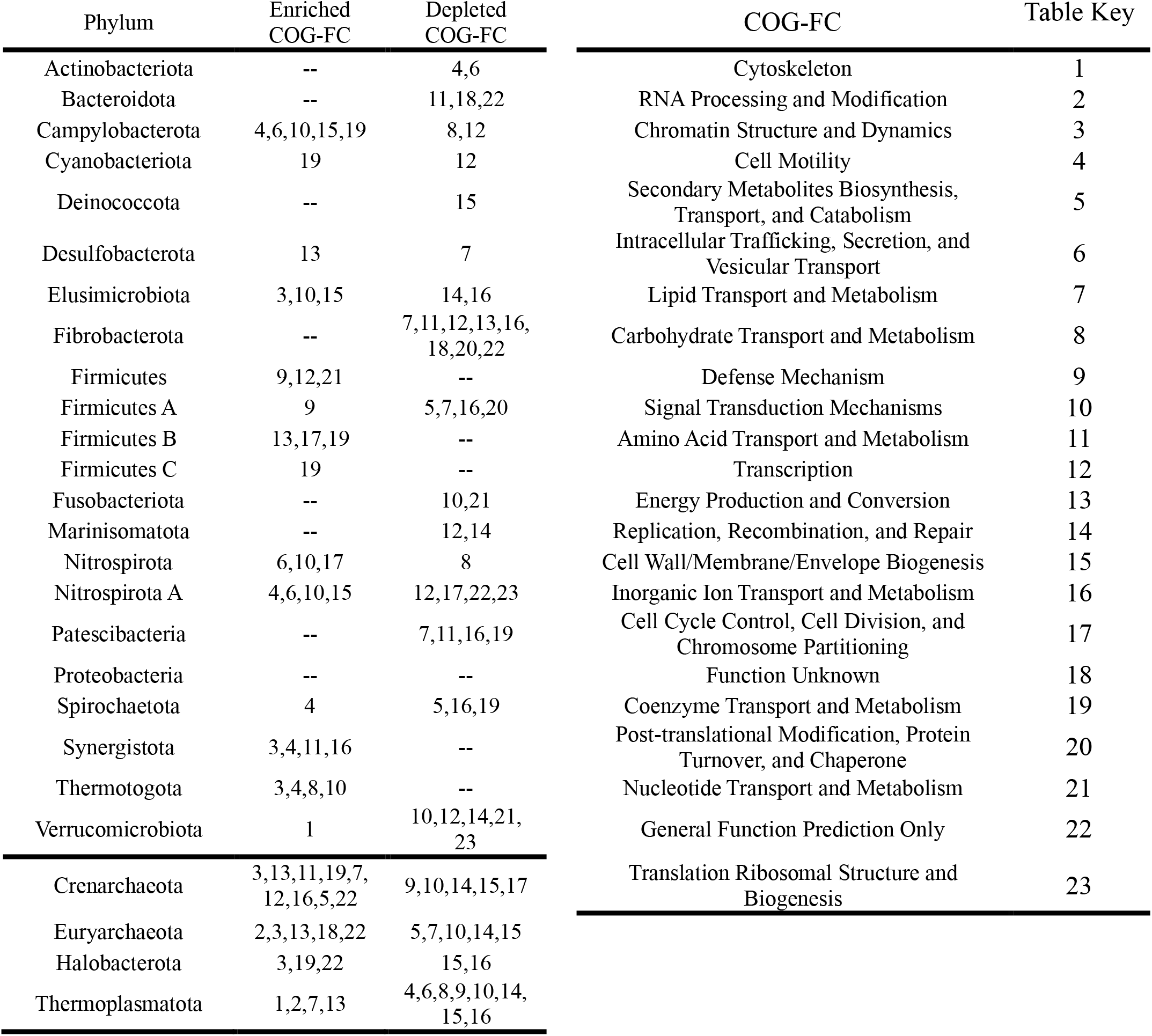
Phylum highly enriched (>85^th^ percentile) or depleted (<15^th^ percentile) in COG-FCs and depletion. All reported categories are statistically significant.

## Discussion

We find that the abundance of each COG-FC in genomes scales with the genome size, so PCA analysis of COG-FC enrichments or depletions successfully distinguished phyla only once COG-FC abundance was normalized to the abundance that was predicted based on genome size (Fig 1C, D). The relationship between COG-FC abundance and genome size was clearly nonlinear, however, with higher slopes at lower genome sizes, indicating that a simple linear model would not accurately capture that relationship (Fig S2).

The PERMANOVA (Fig 2) and analysis of diversity of genomic composition within phyla (Fig 3) showed that microbial lineages exhibit characteristic enrichments of COG-FC, and that the extent of variation varies among taxonomic ranks. Of all the taxonomic ranks, phylum was the most powerful predictor of COG-FC enrichment, which is consistent with observations that phylum can be informative of microbial function (e.g., 10–12). Surprisingly, lower taxonomic ranks such as genus and family had little explanatory power. Many studies focus on metabolic coherence of individual traits and regularly find traits conserved on the family level (2, 3, 13). The discrepancy between previous observations and our observation likely relates to how we characterize patterns in metabolic potential. The tradeoff of the approach used here is that, by analyzing COG-FCs, we lose information about specific genes or potential metabolic functions but gain the ability to apply a consistent analysis across an entire genome and across the entire microbial tree of life. Thus, the extent that observed patterns (Fig 1) reflect phenotypically-expressed differences among lineages is unknown. Nonetheless, the statistical robustness of the relationship between all taxonomic ranks and COG-FC patterns suggests that evolutionary processes (e.g., horizontal gene transfer, vertical gene transfer, duplications, deletions, etc.) control the preponderance of different COG-FCs across lineages.

The role that individual evolutionary processes play in influencing COG-FC enrichments at a given taxonomic rank is likely variable. For instance, horizontal gene transfer is more common among more closely related lineages (14) and thus, likely promotes increased levels of similarity at lower taxonomic ranks. At higher taxonomic ranks, vertical processes may be more important. The asymptote in the mean log_10_-distance from the centroid as function of lineages in a phylum suggests that identifying more lineages for more poorly represented lineages should expand the diversity of COG-FCs that are found, whereas phyla that were adequately sampled (at least ~1000 lineages) exhibited comparable variability in COG-FC distributions (Fig 3B). Since many more than ~1000 distinct lineages of each phylum are likely to exist (15), we propose that the taxonomic rank of phylum implies a fairly consistent degree of diversity in COG-FC distribution. To the extent that phenotype matches genotype at the level of COG-FC distributions, therefore, we expect that typical phyla exhibit similar phenotypic diversity. A notable exception is the phylum Crenarchaeota, which were far more diverse than would be expected based on the number of lineages sampled. The Crenarchaeota, as defined in the GTDB, collapsed members of several phyla that had been designated separately under previous taxonomies, including lineages that had previously been assigned as Crenarchaeota, Thaumarchaeota, Euryarchaeota, Verstraetearchaeota, Korarchaeota, and Bathyarchaeota (5). It is possible that the relationship between marker genes used in the GTDB and the rest of the genome is unusual for this clade, compared to other phyla, or that the GTDB classification of Crenarchaeota is lacking in some other way.

Although the ranks, genus and family, explained relatively little of the variance in COG-FC distribution, blocks of consistent colors were evident in Fig 4, indicating that enrichments or depletions of specific COG-FCs were conserved across each taxonomic rank in some parts of the phylogenetic tree. This is explained by ‘distantly’ (i.e., non-sister clades) related clades occupying similar COG-FC trait-space. The coherence in metabolic potential at higher taxonomic ranks may help explain the distribution of microbial clades across ecological niches. Analyses of habitat associations (9, 16) and single traits (1–3) also support this idea. Our analysis provides quantitative evidence to this idea by demonstrating coherence in metabolic potential with broad-scale patterns in genomic data (Fig 1-4). The question remains: how well do the observed COG-FC enrichments reflect expressed functional traits (i.e., phenotype) across these lineages? It is difficult to address this question systematically, but some of the enrichments and depletions here appear consistent with known physiologies of clades. For instance, Rickettsiales were depleted in nucleotide metabolism and transport, consistent with previously observed lack of a metabolic pathway for purine synthesis among five example Rickettsiales (17). Another example is the depletion in the COG-FCs for energy production and conversion, amino acid transport and metabolism, and carbohydrate transport and metabolism within the Bacteroidetes, Spirochaetes, and Chlamydiales clade. This clade is known to contain many host-dependent pathogens and symbionts (18–20), which are often depleted in these COG-FCs (21).

The GTDB classification is the first fully algorithmic and quantitatively self-consistent microbial taxonomy that can be applied across the tree of life (5). By standardizing the meaning of taxonomic ranks, it creates an objective basis on which to compare microbial functionality to phylogeny. The data presented here support the idea that clades in this taxonomy are functionally coherent, as well as the idea that all taxonomic ranks provide information about the metabolic and ecological niches that individual microbes occupy.

## Materials and Methods

### Genome Database Curation

All bacterial and archaeal genomes from the RefSeq database v92 (22), uncultured bacterial and archaeal (UBA) metagenome-assembled genomes (MAGs) reported in Parks et al. (5, 23), bacterial and archaeal MAGs from Integrated Microbial Genomes and Microbiomes (IMG/M), and bacterial and archaeal single amplified genomes (SAGs) from IMG/M were curated into a single database. All genomic content within the curated database is referred to as “genome(s)” for simplicity. Genomes were assigned taxonomy consistent with the Genome Taxonomy Database (GTDB) using the GTDB toolkit (GTDB-Tk) v0.2.1 (5). The GTDB-Tk taxonomic assignments were consistent with reference package GTDB r86. Lineages which did not receive a genus classification, due to the absence of a reference lineage were excluded from analyses. Due to bias in the abundance of strains in specific clades (e.g., *E. coli*), the lowest taxonomic rank considered during our analysis was species. The COG-FC enrichments (see below) were averaged together for all strains within a given species. An exception was made for lineages which shared a genus classification but lacked a species classification. In this scenario, each genome was treated as an independent lineage. Lastly, only genomes belonging to genera with at least ten unique species in the database were retained. This criterion ensured enough data to generate meaningful statistics during our PERMANOVA. The final database is summarized in Table 1. The genus-level phylogenetic tree was generated from concatenated protein sequence alignments published in Parks et al. (5).

### COG Functional Category Identification, Enumeration, and Normalization

Genes were predicted from individual genomes and translated into protein sequences using Prodigal v.2.6.3 (24). The resulting protein sequences were analyzed for COGs (8). COG position-specific scoring matrices (PSSMs) were downloaded from NCBI’s Conserved Domain Database (27 March 2017 definitions). COG PSSMs were BLASTed against protein sequences with the Reverse Position Specific-BLAST (RPS-BLAST) algorithm (25). Following previously a reported protocol (25), we used an E-value cutoff of 0.01 to assign COGs with RPS-BLAST. The retrieved COGs were assigned to their respective COG functional categories (COG-FCs; 25 in total) and the abundance of each functional category was tabulated using cdd2cog (26) for each genome. The abundance for individual COG-FCs was normalized by the respective COG-FC standard deviation across all lineages. For the COG-FCs, extracellular structures and nuclear structures, the standard deviation was 0. Consequently, data could not be normalized, and thus, these two categories were discarded from all analyses.

COG-FC abundances were normalized by their respective regression slopes of COG-FC abundance for a given genome as a function of genome size. COG-FC abundances were modelled as a function of genome size for individual categories using a generalized additive model (GAM) with a smoothing term due to the pairwise response to genome size (Sup. Figure 1). We used the gam function from the R package, mgcv (27). In some instances, regression fits were visibly skewed by high-leverage data points. High-leverage data were filtered using the influence.gam function in the mgcv package. Data in the 99.5% percentile for influence were excluded when performing regression analysis but were included in all downstream analyses. All regressions were significant with *p*<0.001.

### Principal Component Analysis (PCA)

We performed PCA on the normalized COG-FC abundances and enrichments. Prior to PCA, assumptions of normality were achieved by performing a boxcox transformation on individual COG-FC abundance and enrichment distributions with the boxcox function from the R Package, MASS (28). The resulting distributions were then scaled by the respective COG-FC standard deviation calculated from all genomes. PCA was performed using the princomp function from the R package, stats (29).

### Quantifying COG-FC Variance Explained by Taxonomic Rank

We performed permutational multivariate analysis of variance (PERMANOVA) using the adonis function from the R package, vegan (30). The taxonomic ranks domain, phylum, class, order, family, and genus as well as cultured-status were used as test categorical variables for quantifying variance in COG-FC enrichments explained by the mean taxonomic rank centroids. The default, 999 permutations test, was performed using each categorical variable. Distances were calculated between mean phyla COG-FC enrichment centroids and the respective genomes within that phyla by performing an analysis of multivariate homogeneity of groups dispersions with the betadisper function from the R package, vegan (30). The distance matrix used for both the adonis and betadisper analyses was generated calculating Euclidean distance on the normalized COG-FC enrichments.

The mean log_10_-distance from phylum centroid for each phylum and modeled with the following equation, which represents a hyperbola shifted on the x-axis to ensure that mean distance is zero when n = 1:

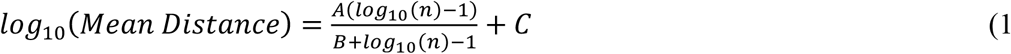

where *A, B*, and *C* are fit coefficients and *n* is the total number of lineages in the given phylum. The Akaike Information Criterion was calculated with the fit from eq 1 using the AIC function from the R package, stats (29).

### Data Availability

The genomes analyzed for the current study are available in NCBI’s RefSeq database (ftp://ftp.ncbi.nlm.nih.gov/refseq/release/). UBA MAGs used for the current study are available under NCBI’s BioProject PRJNA417962 and PRJNA348753. JGI IMG/M acquired from Chad Burdyshaw. Associated genome accessions are available in Dataset S1.

## Supporting information

Supplemenetal_data_description

## Acknowledgments

Funding for this project was provided by a C-DEBI graduate fellowship to TR and an-kind grant of resources from the University of Tennessee / Oak Ridge National Lab Joint Institute for Computational Sciences (JICS) to ADS. We thank Chad Burdyshaw of JICS for help obtaining the genomes used in this project. This is C-DEBI contribution number [to be assigned upon manuscript acceptance].

