## Supplementary material for "Quantitatively Partitioning Microbial Genomic Traits among Taxonomic Ranks across the Microbial Tree of Life": Supplemenetal_data_description

Dataset S1:

Individual rows correspond to individual genomes (excluding the top row which are column headers). Columns 1 through 25 correspond to raw abundances for each COG functional category. Column 26 corresponds to the total number of COGs in a genome. Columns 27, 28, 29, 30, 31, 32, and 33, correspond to the GTDB domain, phylum, class, order, family, genus, and species classification, respectively. Column 34 corresponds to the culture-status. Column 35 is the genomes size in base pairs. Column 36 corresponds to the accession number for each genome. Accessions starting with GCF and GCA are from Refseq and Genbank, respectively. Accessions that are numbers only correspond to IMG/G. Column 37 corresponds to the total number of open reading frames in the genome.

Dataset S2

Individual rows correspond to individual genus-level lineages (excluding the top row which are column headers). Columns 1, 2, 3, 4, and 5 correspond to domain, phylum, order, family, and genus, respectively. Columns 6 through 28 correspond to average enrichments for the respective lineage and COG functional category.
